# ChAdOx1 NiV vaccination protects against lethal Nipah Bangladesh virus infection in African green monkeys

**DOI:** 10.1101/2021.07.20.452991

**Authors:** Neeltje van Doremalen, Victoria A. Avanzato, Friederike Feldmann, Jonathan E. Schulz, Elaine Haddock, Atsushi Okumura, Jamie Lovaglio, Patrick W. Hanley, Kathleen Cordova, Greg Saturday, Teresa Lambe, Sarah C. Gilbert, Vincent J. Munster

## Abstract

Nipah virus (NiV) is a highly pathogenic and re-emerging virus which causes sporadic but severe infections in humans. Currently, no vaccines against NiV have been approved. We previously showed that ChAdOx1 NiV provides full protection against a lethal challenge with NiV Bangladesh (NiV-B) in hamsters. Here, we investigated the efficacy of ChAdOx1 NiV in the lethal African green monkeys (AGMs) NiV challenge model. AGMs were vaccinated either 4 weeks before challenge (prime vaccination), or 8 and 4 weeks before challenge with ChAdOx1 NiV (prime-boost vaccination). A robust humoral and cellular response was detected starting 14 days post initial vaccination. Upon challenge, control animals displayed a variety of signs and had to be euthanized between 5- and 7-days post inoculation. In contrast, vaccinated animals showed no signs of disease, and we were unable to detect infectious virus in all but one swab and all tissues. Importantly, no to limited antibodies against fusion protein or nucleoprotein IgG could be detected 42 days post challenge, suggesting that vaccination induced a very robust protective immune response preventing extensive virus replication.

**One Sentence Summary:** A single vaccination with ChAdOx1 NiV protects African green monkeys against lethal disease induced by Nipah virus inoculation.

## Introduction

Nipah virus (NiV) is a highly pathogenic re-emerging member of the *Paramyxovirus* family, genus *Henipavirus*. NiV causes sporadic infections in humans, resulting in severe neurological and respiratory disease, often with a fatal outcome. NiV was first detected in 1998, when the strain NiV-Malaysia (NiV-M) caused an outbreak of severe encephalitis in pig farmers from Malaysia and Singapore, with a case-fatality rate of 38%^1^. Since 2001, outbreaks with a closely related strain, NiV-Bangladesh (NiV-B), have occurred almost yearly in Bangladesh^2^, resulting in 319 accumulated cases and 225 associated deaths (case-fatality rate 71%)^3^. Most recently, outbreaks have also been reported in India^4^.

The natural reservoir of NiV is the *Pteropus spp*. fruit bat^5–7^. Outbreaks in Bangladesh and India have been associated with the consumption of date palm sap, which may have been contaminated with bat urine^8–10^. In contrast, in Malaysia and Singapore, pigs were the intermediate host, likely infected via the consumption of mango fruits, which were contaminated with NiV after partial consumption by bats^11^. Importantly, human-to-human transmission of NiV has also been reported^12,13^.

Although the total number of cases caused by NiV are limited, the virus causes severe disease and transmits between humans^12,13^, and can infect a wide range of animals^14^, and thus NiV is categorized by the WHO as a pathogen with epidemic potential which poses a great public health risk and requires research aimed at the development of countermeasures.

Several vaccine candidates have been evaluated in animal models^15^. The most extensively studied vaccine to date is based on the glycoprotein of Hendra virus (HeV), another member of the genus *Henipavirus*^16^. HeV-sG, a soluble form of the HeV receptor binding glycoprotein, was marketed by Zoetis, Inc., in Australia as an equine vaccine against HeV under the name Equivac^®^ HeV. It is the first commercialized vaccine against a BSL-4 agent^17^.

Recently, it was shown that HeV-sG vaccination can protect African green monkeys (AGMs) against lethal NiV disease as early as 7 days post immunization^17^. Enrollment has started for a Phase I randomized placebo-controlled clinical trial on NiV-vaccine candidate HeV-sG-V, which is based on HeV-sG, with results expected in October 2021 (ClinicalTrials.gov NCT04199169). This clinical trial is the first of its kind for NiV and is the result of a global partnership between Auro Vaccines LLC and the Coalition of Epidemic Preparedness Innovations (CEPI).

In the current study, we are testing efficacy of a different NiV vaccine candidate in AGMs. ChAdOx1 is a replication-deficient simian adenoviral vector, which has been developed for a multitude of different pathogens by the University of Oxford. A vaccine based on this vector named ChAdOx1 nCoV-19 (also known as AZD1222, Vaxzevria, or Covishield) has been developed against severe acute respiratory syndrome coronavirus 2 (SARS-CoV-2), the etiological agent of COVID-19. ChAdOx1 nCoV-19 has been fully approved in Brazil and approved for emergency use in 64 additional countries. The effectiveness of the vaccine is 92% and is 79% against symptomatic infection^18,19^.

We have previously demonstrated that a single dose of ChAdOx1 NiV, which encodes the receptor binding protein (G) of NiV-B, fully protected Syrian hamsters against a lethal dose of NiV-B or NiV-M^20^. In the current study, we investigate whether ChAdOx1 NiV is protective in the lethal NiV AGM model^21^. We show that a single dose of ChAdOx1 NiV results in a robust innate and adaptive immune response, which is fully protective against lethal disease in AGMs. Furthermore, no to limited immune response against nucleoprotein or fusion protein was detected in vaccinated animals post challenge, but was detected in control challenged animals, suggesting that vaccination provides near complete protection against NiV infection.

## Results

### ChAdOx1 NiV vaccination of African green monkeys elicits a potent adaptive immune response

Four animals per group were vaccinated via the intramuscular (I.M.) route with ChAdOx1 NiV using a prime-boost regimen at 56 and 28 days before challenge, or a prime-only regimen at 28 days before challenge. As a control, four animals were vaccinated via the I.M. route with ChAdOx1 GFP at 56 and 28 days before challenge. Binding antibody titers against NiV G protein were determined at day of vaccination, 14 days post vaccination, and day of challenge. NiV G-specific IgG antibodies could be detected as early as 14 days post vaccination and were significantly increased upon boost vaccination (Two-tailed Mann-Whitney test, p=0.0286). All vaccinated animals had detectable NiV G-specific antibodies on the day of challenge (Figure 1A). No NiV G-specific IgG antibodies were detected in control animals. Likewise, virus neutralizing antibodies were detected in all vaccinated animals at day of challenge and were significantly increased upon boost vaccination (Figure 1B, Mann-Whitney test, p=0.0286). NiV G-specific T cell responses were investigated using a peptide library divided into six peptide pools that spanned the full length of NiV G. A single vaccination resulted in specific T cell responses in all animals, and a subsequent boost vaccination raised T cell responses significantly above control T cell responses (Figure 1C, Mann-Whitney test, p = 0.0271).

**Figure 1.**
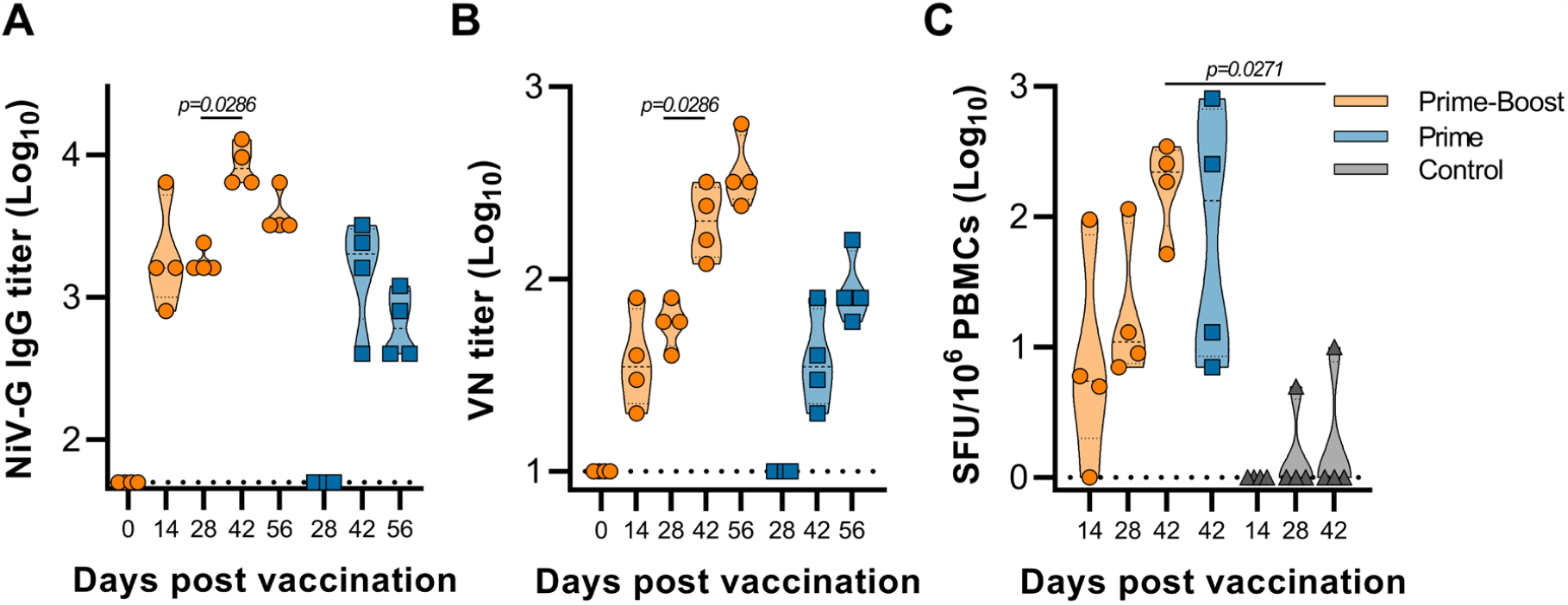
Vaccination with ChAdOx1 NiV in AGMs induces humoral and cellular immune responses. **(A)** Truncated violin plot shows evidence of NiV-G-specific IgG in serum at the indicated days post prime vaccination in animals receiving intramuscular ChAdOx1 NiV via a prime-boost (orange, n=4) or prime-only regimen (blue, n=4). **(B)** Truncated violin plot of neutralizing antibodies in serum are shown. **(C)** Truncated violin plot of NiV-G protein-specific T cell responses in PBMCs isolated from vaccinated or controls animals at indicated time points minus 0 days post vaccination (DPV) response. SFU, spot-forming units. Black lines indicate median; dotted lines indicate quartiles. The dotted line shows the limit of detection.

### Vaccinated animals do not show signs of disease

Animals were inoculated with 2 ⨯ 10^5^ TCID_50_ of NiV-B, split equally between the intranasal and intratracheal route. Animals were checked daily for clinical signs. Animals vaccinated with ChAdOx1 GFP displayed a variety of signs starting at 3 days post inoculation (DPI), including neurological signs and respiratory signs (Table S1). All four animals in this group reached an endpoint clinical score of 35 or higher between 5 to 7 DPI and were euthanized. In contrast, no signs of disease were observed in animals vaccinated with ChAdOx1 NiV (Figure 2A-B, Table S1). Exams were performed on 0, 3, 5, 7, 10, 14, 21, 28, 35, and 42 DPI. Radiographs were scored as previously described^22^. Whereas no to limited changes from baseline were observed in ChAdOx1 NiV vaccinated animals (score between 0-2), radiograph scores of control-vaccinated animals started increasing at 3 DPI (score between 0-3) and continued to increase until day of necropsy (score between 7-14, Figure 2C). Throat and nose swabs were collected on all exam days and the presence of infectious virus was investigated. Infectious virus could be detected in both nose and throat swabs of 3 out of 4 control-vaccinated animals. In contrast, all swabs obtained from animals vaccinated with ChAdOx1 NiV were negative, except for one throat swab obtained at 3 DPI from one animal (animal 1) in the prime-boost group (Figure 2D-E). The presence of binding antibodies against fusion glycoprotein (F) and nucleoprotein (N) of NiV was then investigated in sera obtained from vaccinated animals at 42 DPI. A low titer of binding antibodies against N was found in sera from animal 1, and antibodies against F were detected in sera from animal 1, 7, and 8 (Figure 2F).

**Figure 2.**
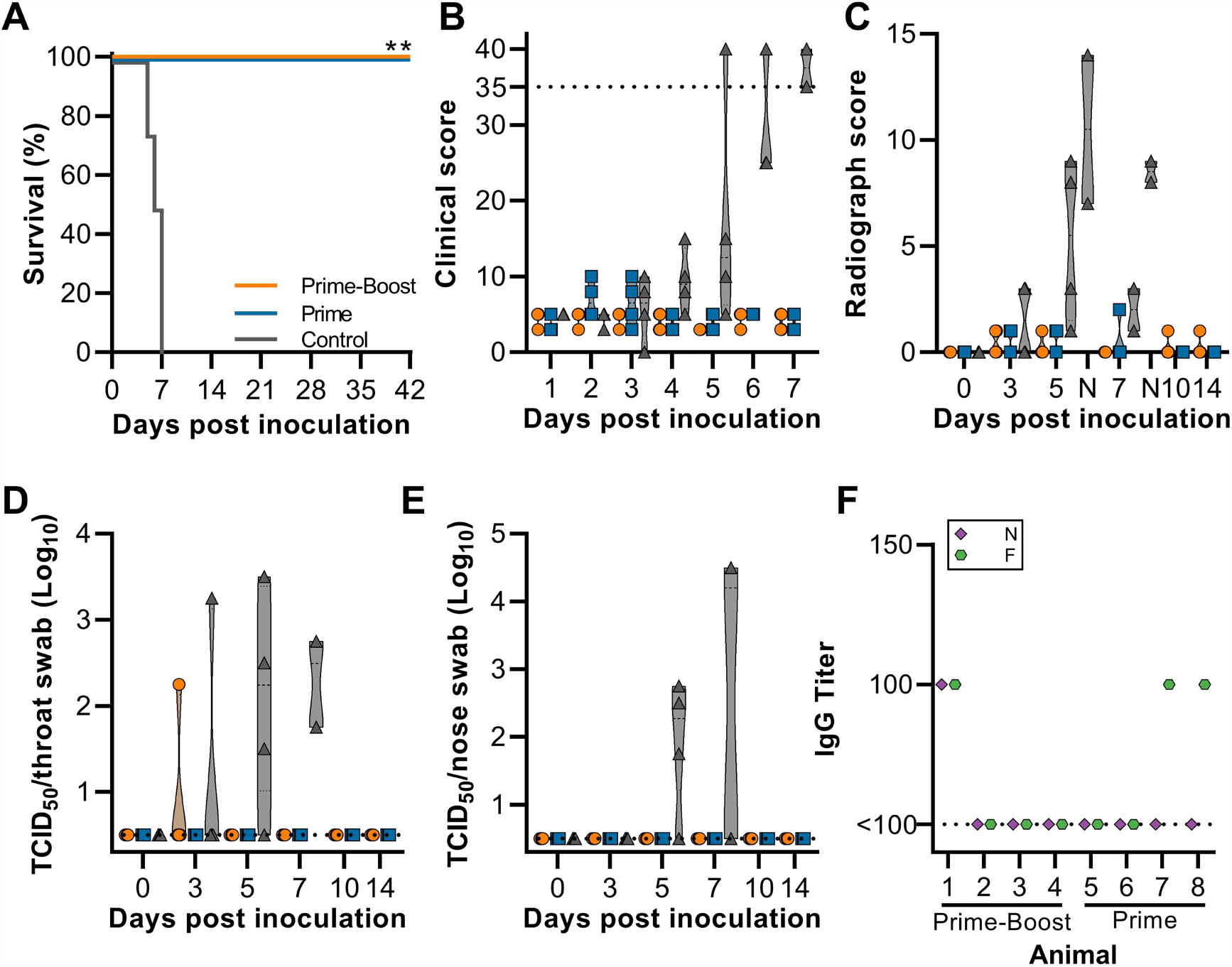
Clinical signs and NiV detection in vaccinated AGMs upon virus challenge. 28 days post final vaccination, AGMs (N=4 per group) were inoculation with NiV-B and monitored for up to 42 days. **(A)** Survival of AGMs. **(B)** Truncated violin plot of daily clinical score. Dotted line indicates score requiring euthanasia. **(C)** Truncated violin plot of thoracic radiograph scores on exam days and day of necropsy (N, controls only). **(D/E)** Truncated violin plots of infectious NiV in throat swabs (D) and nose swabs (E). At 7 DPI, only 2 control animals were part of the study. Dotted line indicates limit of detection. **(F)** Truncated violin plot shows lack of evidence of NiV-nucleoprotein (N, purple diamond) or NiV-fusion glycoprotein (F, green octagon) - specific IgG in serum at 42 DPI. Dotted line indicates limit of detection. For all panels, orange indicates prime-boost vaccinated animals, blue indicates prime only vaccinated animals, and grey indicates control animals.

### No evidence of NiV detected in tissues of vaccinated animals

Animals were euthanized when euthanasia criteria were reached (control group, 5-7 DPI) or at 42 DPI (end of study, vaccinated animals). Lung:body weight ratio, indicative of edema, was higher for control animals than for vaccinated animals (Figure 3A). At necropsy, the percentage of each lung lobe that showed lesions was scored. Whereas a high percentage of gross lesions was observed in control animals (50 out of 56 lung lobes, median 50%), this was almost completely absent in vaccinated animals (9 out of 56 lung lobes, median 0% for both prime-boost and prime-only vaccinated groups) (Figure 3B). Lungs of control animals failed to collapse (3 out of 4), and pleural effusion was observed in all animals. Cervical lymph node enlargement and edematous mediastinal lymph nodes were observed in 2 and 3 out of 4 control animals, respectively. One animal showed a diffuse hemorrhage from the medulla oblongata to the cervical spinal cord, with petechiae on the cerebellum.

**Figure 3.**
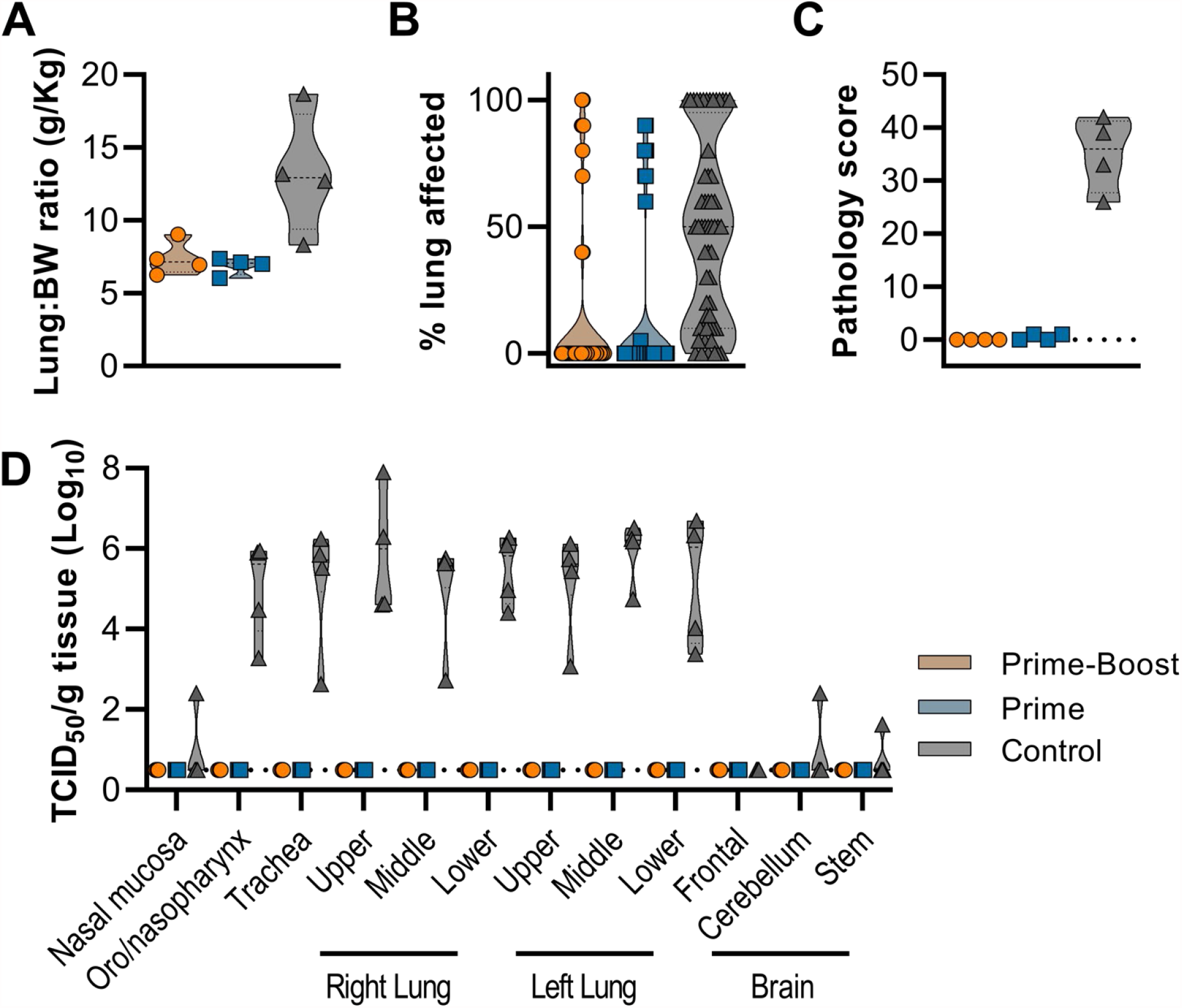
No evidence of NiV infection detected in vaccinated animals. AGMs were euthanized when a clinical score of 35 was reached (all control animals, 5-7 DPI) or on 42 DPI (all vaccinated animals). **(A)** Truncated violin plot of lung:body weight (BW) ratio. **(B)** Truncated violin plot of gross lung lesions. which were scored for each lung lobe, ventral and dorsal. **(C)** Truncated violin plot of pathology score. All six lung lobes were scored and cumulative score is shown per animal (maximum score = 90). **(D)** Truncated violin plot of infectious virus detected in respiratory tract and brain tissue. For all panels, orange indicates prime-boost vaccinated animals, blue indicates prime only vaccinated animals, and grey indicates control animals. No statistical tests were performed since samples were obtained on different days post challenge.

Histologically, no pulmonary pathology consistent with NiV lesions was observed in vaccinated animals (Figure 4A-B), and subsequently the pathology score was low (Figure 3C). Likewise, no NiV RNA staining was detected in lung tissue (Figure 4D-E). In stark contrast, histological lesions were present in lung tissue obtained from all control animals and were characterized as multifocal, random, minimal (1-10%) to marked (51-75%) bronchointerstitial pneumonia. The pneumonia was characterized by thickening of alveolar septa by edema fluid and fibrin and small to moderate numbers of macrophages, syncytial cells and neutrophils (Figure 4C). In situ hybridization reveals abundant viral RNA distributed throughout lesions in vascular endothelium and type I&II pneumocytes in tissue from control animals (Figure 4F). Infectious virus was only detected in tissue obtained from control animals, and not in tissue obtained from vaccinated animals (Figure 3D, S1).

**Figure 4.**
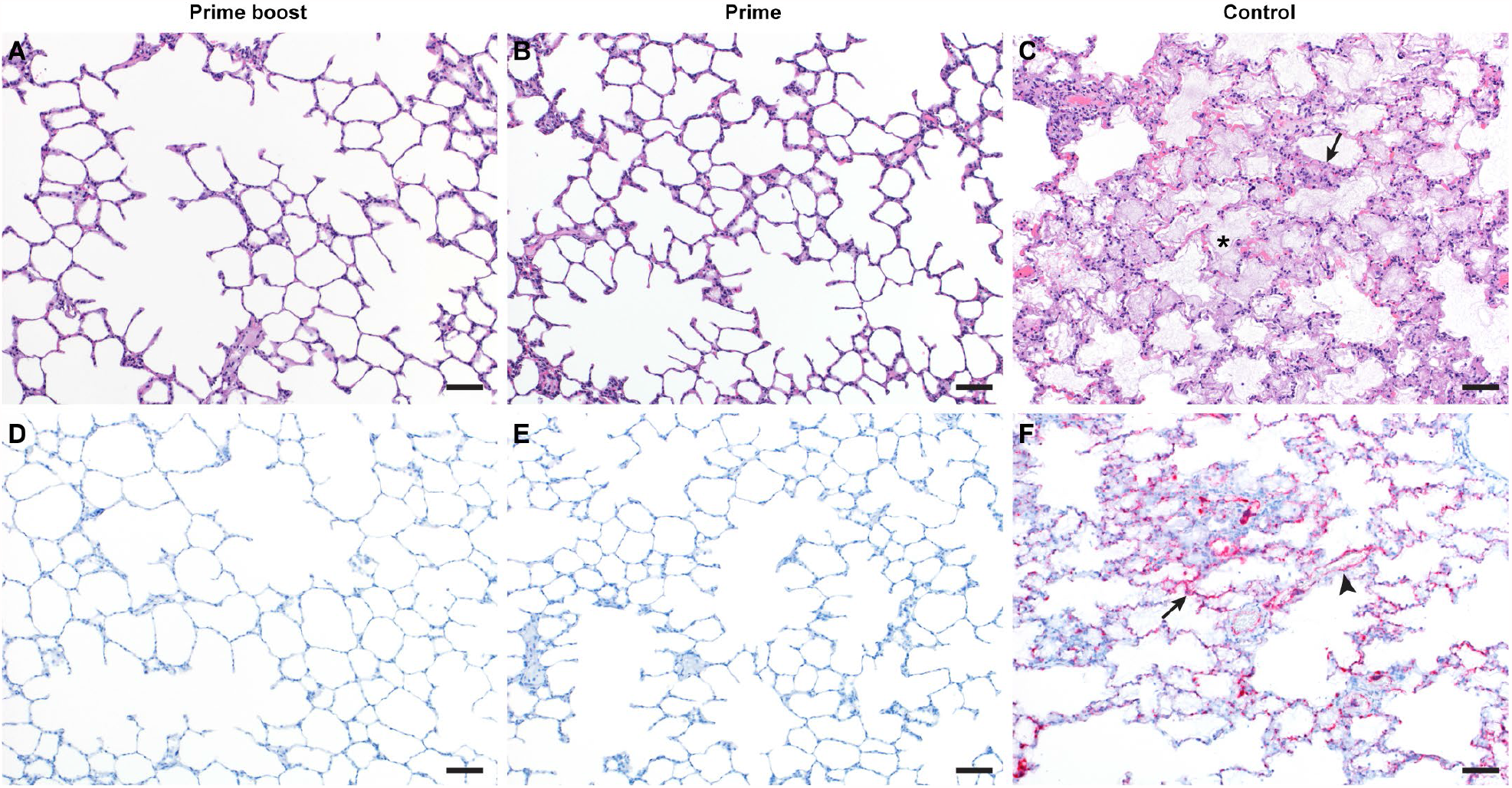
Pulmonary effects of ChAdOx1-NiV vaccine efficacy in the AGM model of NiV infection. **(A-C)** Lung tissue sections were stained with hematoxylin and eosin. (A-B) no pathology was observed. (C) moderate to marked interstitial pneumonia (arrow) with abundant fibrin and edema (asterisk). **(D-F)** In situ hybridization for NiV RNA, resulting in a red stain. (D-E) no immunoreactivity was observed. (F) Immunoreactive vascular endothelium (arrowhead) and pneumocytes (arrow). 100x bar = 50µm.

## Discussion

We evaluated the vaccine candidate ChAdOx1 NiV, which we previously showed to be fully protective in Syrian hamsters^20^, in the lethal AGM NiV challenge model. We show here that a single dose of ChAdOx1 NiV was fully protective against a lethal challenge with NiV-B in AGMs. Furthermore, we found very limited evidence of virus replication in vaccinated animals; all but one swab was negative for infectious virus, no virus was found in tissues obtained from vaccinated animals, and no to a very limited immune response was detected against NiV F or N proteins after challenge of vaccinated animals. These data suggest the vaccine may provide close to complete protective immunity in AGMs.

Bossart *et al*.^23^ show a similar lack of NiV-F specific antibodies in vaccinated animals. In that study, AGMs were vaccinated with a HeV-G subunit vaccine 6 and 3 weeks before challenge with a lethal dose of NiV. Serum was obtained throughout the experiment and NiV F-specific IgM could not be detected at any point. Furthermore, whereas vaccination induced high IgG titers against NiV-G and HeV-G, there was no increase in G-specific IgG titers after challenge with NiV. Together, these data suggest a lack of virus replication in vaccinated animals^23^. Likewise, in a study performed by Lo *et al*.^24^, hamsters were vaccinated with replication-deficient VSV-based NiV G or NiV F. VN titers were measured in serum obtained at 28 days post vaccination and 32 days post challenge. No anamnestic response was detected in vaccinated animals, suggesting that this vaccine may also provide (near) complete proimmunity^24^. In a study by Prescott *et al*., AGMs were vaccinated with rVSV-EBOV-GP-NiVG, 29 days before challenge with NiV Malaysia. An increase in binding and neutralizing antibodies titer between 0 DPI and 16/17 DPI (termination of study) was found in 2/3 vaccinated animals^25^.

The approval of a NiV vaccine is hindered by the feasibility of an efficacy trial due to the sporadic nature of infections, the large geographical area where the spillover occurs and the low number of annual cases^26^. To address these complications, the U.S. Food and Drug Administration implemented the ‘Animal Rule’ in 2002^27^. This rule can be utilized to establish efficacy based on studies performed in animal models that faithfully recapitulate human disease, such as the AGM and hamster NiV model^26^. Thus far, 18 products have been approved via this route^28^. In both hamster and AGM NiV models, vaccination with ChAdOx1 NiV resulted in induction of high antibody titers coupled with complete protection against lethal NiV disease. ChAdOx1 NiV is based on the same vector as ChAdOx1 nCoV-19, which has been approved for emergency use in over 60 countries worldwide. Subsequently, 100 million people have been vaccinated with the vector. Safety profiles obtained in ChAdOx1 nCoV-19 clinical studies^29^ combined with efficacy studies in animal models^20^ may provide sufficient information for approval of ChAdOx1 NiV.

Although several NHP studies have successfully investigated the efficacy of NiV vaccines^23,25,30–32^, thus far only one vaccine has advanced into clinical trials. Here, we show that the widely used ChAdOx1 vector can be modified to provide protection against NiV in a lethal NHP model. Previously, similar work investigating the efficacy of a ChAdOx1 MERS vaccine in rhesus macaques^33^ was instrumental in the development of the ChAdOx1 nCoV-19 vaccine^34^. If the next pandemic were to be caused by a member of the genus *Henipavirus*, the current study could be influential in the development of a rapid vaccine. Future studies, such as Phase I clinical trial studies and further efficacy studies in AGMs, should aim to obtain approval via the FDA Animal Rule.

## Materials and Methods

### Ethics Statement

The Institutional Animal Care and Use Committee (IACUC) at Rocky Mountain Laboratories approved all animal study requests, which were conducted in an Association for Assessment and Accreditation of Laboratory Animal Care (AAALAC)-accredited facility, following the basic principles and guidelines in the Guide for the Care and Use of Laboratory Animals 8th edition, the Animal Welfare Act, United States Department of Agriculture and the United States Public Health Service Policy on Humane Care and Use of Laboratory Animals.

Animals were kept in climate-controlled rooms with a fixed light/dark cycle (12-hours/12-hours). African green monkeys were housed in individual primate cages allowing social interactions, fed a commercial monkey chow, treats and fruit with ad libitum water and were monitored at least twice daily.

Environmental enrichment consisted of a variety of human interaction, commercial toys, videos, and music. The Institutional Biosafety Committee (IBC) approved work with infectious Nipah virus strains under BSL4 conditions. All sample inactivation was performed according to IBC approved standard operating procedures for removal of specimens from high containment^35,36^.

### Study design animal experiments

12 African green monkeys (6M, 6F) between 3-5 years old were sorted by sex, then by weight, and then randomly divided into three groups of four animals. Group 1 was vaccinated with ChAdOx1 NiV at -56 and -28 DPI, group 2 was vaccinated with ChAdOx1 NiV at -28 DPI, group 3 was vaccinated with ChAdOx1 GFP at -56 and -28 DPI. All vaccinations were done intramuscularly with 2.5 ⨯ 10^10^ VP/animal diluted in sterile PBS. Animals were challenged with Nipah Bangladesh (AY988601) diluted in sterile DMEM at 0 DPI; 4 mL intratracheally (2.5 ⨯ 10^4^ TCID_50_/mL) and 1 mL intranasally (1 ⨯ 10^5^ TCID_50_/mL). Animals were scored daily by the same person who was blinded to study group allocations using a standardized scoring sheet^37^. Scoring was based on the following criteria: general appearance, skin and coat appearance, discharge, respiration, feces and urine appearance, appetite, and activity. Clinical exams were performed on -56, -42, -28, -14, 0, 3, 5, 7, 10, 14, 21, 28, 35, and 42 DPI. Blood, nasal, and throat swabs were collected on all exam dates. Hematology analysis was completed on a ProCyte DX (IDEXX Laboratories, Westbrook, ME, USA) and the following parameters were evaluated: red blood cells (RBC), hemoglobin (Hb), hematocrit (HCT), mean corpuscular volume (MCV), mean corpuscular hemoglobin (MCH), mean corpuscular hemoglobin concentration (MCHC), red cell distribution weight (RDW), platelets, mean platelet volume (MPV), white blood cells (WBC), neutrophil count (abs and %), lymphocyte count (abs and %), monocyte count (abs and %), eosinophil count (abs and %), and basophil count (abs and %). Serum chemistries were completed on a VetScan VS2 Chemistry Analyzer (Abaxis, Union City, CA) and the following parameters were evaluated: glucose, blood urea nitrogen (BUN), creatinine, calcium, albumin, total protein, alanine aminotransferase (ALT), aspartate aminotransferase (AST), alkaline phosphatase (ALP), total bilirubin, globulin, sodium, potassium, chloride, and total carbon dioxide. Ventro-dorsal and right/left lateral radiographs were taken on clinical exam days prior to any other procedures. Radiographs were evaluated and scored for the presence of pulmonary infiltrates by two board-certified clinical veterinarians according to a standard scoring system^22^. Briefly, each lung lobe (upper left, middle left, lower left, upper right, middle right, lower right) was scored individually based on the following criteria: 0 = normal examination; 1 = mild interstitial pulmonary infiltrates; 2 = moderate interstitial pulmonary infiltrates, perhaps with partial cardiac border effacement and small areas of pulmonary consolidation (alveolar patterns and air bronchograms); and 3 = pulmonary consolidation as the primary lung pathology, seen as a progression from grade 2 lung pathology. Day 0 radiographs are taken prior to inoculation, and thus serve as a baseline for each animal. All subsequent radiographs are compared to the Day 0 radiographs, evaluated for changes from baseline and scored based on the criteria noted above. At study completion, thoracic radiograph findings are reported as a single cumulative radiograph score for each animal on each exam day; scores may range from 0 to 18. Necropsy was performed on 42 DPI or when euthanasia criteria was reached.

### Generation of vaccine ChAdOx1 NiV

ChAdOx1 NiV was produced as previously described^20^. Briefly, the G gene from Nipah virus (Bangladesh outbreak 2008-2010, Genbank accession number: JN808864.1) was codon optimized for humans, synthesized by GeneArt (Thermo Fisher Scientific), and cloned into a transgene expression plasmid comprising a modified human cytomegalovirus immediate early promoter (CMV promoter) with tetracycline operator (TetO) sites and the polyadenylation signal from bovine growth hormone (BGH). This expression cassette was inserted into the E1 locus of the genomic clone of ChAdOx1 using site-specific recombination^38^. The virus was rescued and propagated in T-REx-293 cells (Invitrogen). Purification was by CsCl gradient ultracentrifugation, virus titers were determined by hexon immunostaining assay and viral particles were calculated based on spectrophotometry^39,40^.

### Cells and virus

NiV (strain Bangladesh/200401066) was kindly provided by the Special Pathogens Branch of the Centers for Disease Control and Prevention, Atlanta, Georgia, United States. This isolate was obtained from a throat swab collected from patient #3001 (10-year old male) in Bangladesh on January 22, 2004. The patient developed altered mental status, cough, and breathing difficulties on January 21. The patient was admitted to Goalando Hospital, Bangladesh, on January 22. None of the patient’s contacts developed NiV infection; the patient is presumed to have been infected via direct spillover from the bat reservoir (Dr. Steve Luby, personal communication)^41^. All virus propagation was performed in VeroE6 cells cultured in Dulbecco’s modified Eagle’s medium (DMEM, Sigma) supplemented with 2% fetal bovine serum (Gibco), 1 mM L-glutamine (Gibco), 50 U/ml penicillin (Gibco), and 50 μg/ml streptomycin (Gibco) (2% DMEM). VeroE6 cells were maintained in DMEM supplemented with 10% fetal calf serum, 1 mM L glutamine, 50 U/ml penicillin and 50 μg/ml streptomycin.

### Titration assay

Virus titrations were performed by end-point titration in VeroE6 cells, which were inoculated with tenfold serial dilutions of virus swab media or tissue homogenates. After 1hr incubation at 37°C and 5% CO_2_, tissue homogenate dilutions were removed, washed twice with PBS and replaced with 100 μl 2% DMEM. Cytopathic effect was scored at 5 DPI and the TCID_50_ was calculated from a minimum of 4 replicates by the Spearman-Karber method^42^.

### Virus neutralization assay

Heat-inactivated sera (30 min, 56 °C) was serially diluted (2x) in 2% DMEM. Hereafter, 100 TCID_50_ of NiV was added. After 1hr of incubation at 37 °C and 5% CO_2_, serum:virus mixture was added to VeroE6 cells and incubated at 37°C and 5% CO_2_. At 5 DPI, cytopathic effect was scored. The virus neutralization titer was expressed as the reciprocal value of the highest dilution of the serum which still inhibited virus replication.

### Production NiV G and F proteins

Nipah proteins were produced as previously described^20^. Briefly, NiV-G Malaysia (residues E144 - T602, gene accession number NC_002728) was cloned into the pHLSEC mammalian expression vector^43^ and NiV-F Malaysia (residues G26 - D482, gene accession number AY816748.1) was cloned into the pHLSEC vector containing a C-terminal GCNt trimerization motif^44^. The constructs were transiently expressed in human embryonic kidney (HEK) 293T cells. Supernatant was diafiltrated using the AKTA FLUX system (GE Healthcare) against either PBS, pH 7.4 (NiV-G) or buffer containing 10 mM Tris and 150 mM NaCl, pH 8.0 (NiV-F). The proteins were further purified by Ni-NTA immobilized metal-affinity chromatography using His-Trap HP columns (GE Healthcare) followed by size exclusion chromatography. NiV-G was purified using a Superdex 200 10/300 Increase GL column (GE healthcare) equilibrated in PBS pH 7.4 and NiV-F was purified using a Superose 6 Increase 10/300 GL column (GE healthcare) equilibrated in 10 mM Tris and 150 mM NaCl pH 8.0.

### Enzyme-linked immunosorbent assay for Nipah G, N, and F proteins

Maxisorp plates (Nunc) were coated overnight at 4°C with 5 µg of G, N (Native Antigen Company) or F protein per plate in Carb/Bicarb binding buffer (4.41 g KHCO_3_ and 0.75 g Na_2_CO_3_ in 1 L distilled water). After blocking with 5% milk in PBS with 0.01% tween (PBST), serum in 5% milk in PBST was incubated at RT for 1 hr. Antibodies were detected using affinity-purified antibody peroxidase-labeled goat-anti-monkey IgG (Seracare) in 5% milk in PBST and TMB 2-component peroxidase substrate (Seracare) and read at 450 nm. All wells were washed 3x with PBST in between steps.

### ELISpot assay and ICS analysis

PBMCs were isolated from ethylene diamine tetraaceticacid (EDTA) whole blood using LeucosepTM tubes (Greiner Bio-one International GmbH) and Histopaque®-1077 density gradient cell separation medium (Sigma-Aldrich) according to the manufacturers’ instructions. The ImmunoSpot® Human IFN-*γ* Single-Color Enzymatic ELISpot Assay Kit was utilized according to the manufacturer’s protocol (Cellular Technology Limited). PBMCs were plated at a concentration of 300,000 cells per well and were stimulated with six contiguous peptide pools spanning the length of the G protein sequence at a concentration of 2 µg/mL per peptide. One peptide (sequence AFNTVIALLGSIVII) was excluded due to false positive results. Analysis was performed using the CTL ImmunoSpot® Analyzer and ImmunoSpot® Software (Cellular Technology Limited). Spot forming units (SFU) per 1.0⨯ 10^6^ PBMCs were summed across the 6 peptide pools for each animal.

### Histology and in situ hybridization

Harvested tissues were fixed for a minimum of 7 days in 10% neutral-buffered formalin and subsequently embedded in paraffin. Hematoxylin and eosin (H&E) staining and in situ hybridization (ISH) were performed on tissue sections and cell blocks. Detection of NiV viral RNA was performed using the RNAscope FFPE assay (Advanced Cell Diagnostics Inc., Newark, USA) as previously described^45^ and in accordance with the manufacturer’s instructions. Briefly, tissue sections were deparaffinized and pretreated with heat and protease before hybridization with target-specific probes for NiV or HeV. Ubiquitin C and the bacterial gene, dapB, were used as positive and negative controls, respectively. Whole-tissue sections for selected cases were stained for NiV and HeV viral RNA, UBC and dapB by the RNAscope VS FFPE assay (RNAscopeVS, Newark, USA) using the Ventana Discovery XT slide autostaining system (Ventana Medical Systems Inc., Tucson, USA). A board-certified veterinary anatomic pathologist blinded to the study groups evaluated all tissue slides. Pathology score was determined by scoring 6 lung lobes for each animal for the following characteristics: lymphoid cuffing; pneumonia, bronchointerstitial, with fibrin, edema and syncytial cells; and perivascular and alveolar edema and fibrin. Scoring was as follows: 0 = no lesions; 1 = 1-10%; 2 = 11-25%; 3 = 26-50%; 4 = 51-75%; 5 = 76-100%. All scores per animal were added to allow a maximum score of 90.

### Statistical analysis

Kruskal-Wallis test for multiple comparisons was used to test for statistical significance. P-values < 0.05 were significant.

## Supplementary Materials

**Figure S1.**
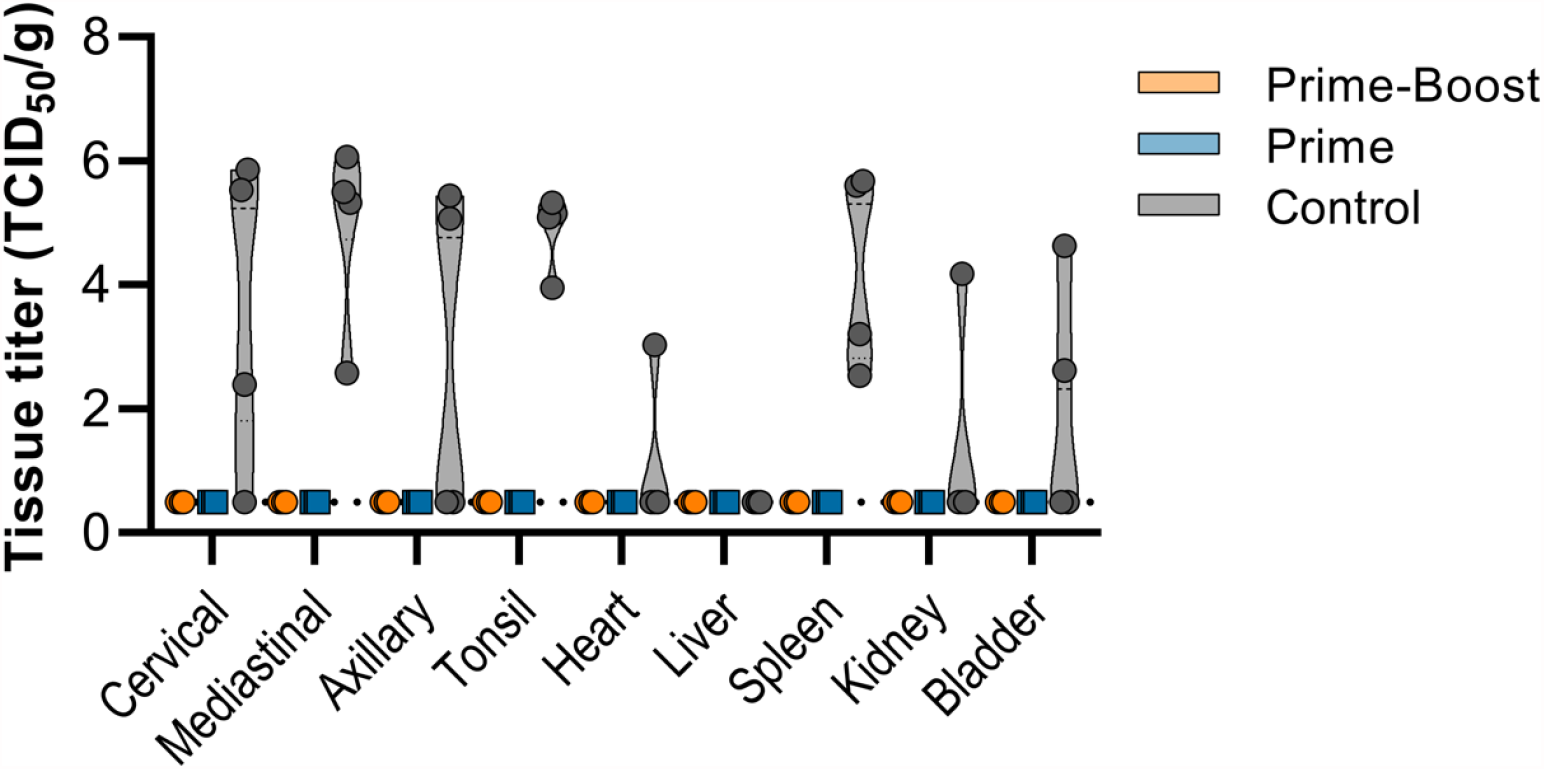
Presence of infectious virus in non-respiratory or brain tissue of African green monkeys inoculated with Nipah virus. Violin plot of infectious virus detected in respiratory tract and brain tissue. Orange circles, prime-boost vaccine; blue squares, prime–only vaccine; grey triangles, controls. No statistical tests were performed since samples were obtained on different days.

**Table S1.** Clinical signs in AGMs inoculated with NiV-B. RA = reduced appetite; RF = ruffled fur; HP = hunched posture; ND= nasal discharge; IR= increased respirations, OM= open mouth breathing, CY= cyanotic, NS= neurological symptoms

| Treatment | Animal | 1 | 2 | 3 | 4 | 5 | 6 | 7 |
| --- | --- | --- | --- | --- | --- | --- | --- | --- |
| Prime-Boost<br>ChAdOx1<br>NiV | 1 | RA, 3 | RA, 3 | RA, 3 | RA, 3 | RA, 3 | RA, 5 | RA, 5 |
|  | 2 | RA, 5 | RA, 5 | RA, 5 | RA, 5 | RA, 3 | RA, 5 | RA, 3 |
|  | 3 | RA, 5 | RA, 5 | RA, 5 | RA, 5 | RA, 3 | RA, 5 | RA, 5 |
|  | 4 | RA, 5 | RA, 5 | RA, 5 | RA, 3 | RA, 3 | RA, 3 | RA, 3 |
| Prime<br>ChAdOx1<br>NiV | 5 | RA, 3 | RA, 8 | RA, 8 | RA, 3 | RA, 3 | RA, 5 | RA, 5 |
|  | 6 | RA, 3 | RA, 10 | RA, 10 | RA, 5 | RA, 5 | RA, 5 | RA, 3 |
|  | 7 | RA, 3 | RA, 5 | RA, 3 | RA, 5 | RA, 3 | RA, 5 | RA, 5 |
|  | 8 | RA, 5 | RA, 5 | RA, 5 | RA, 3 | RA, 5 | RA, 5 | RA, 5 |
| Prime-Boost<br>ChAdOx1<br>GFP | 9 | RA, RF, 3 | RA, RF, 5 | RA, RF, 3 | RA, RF, 8 | RA, RF, 5 | RA, HP, RF, ND, IR 25 | RA, HP, RF, 40<br>Fever 40.3°C, ND, IR, OM, CY |
|  | 10 | RA,5 | RA, 5 | -, 0 | RA, IR, 8 | IR,5 | RA, RF, IR, NS, 25 | HP, RF, RA, IR, NS, 35 |
|  | 11 | RA, 5 | RA, 5 | RA, 5 | RA, IR,10 | HP, RA, IR,NS ,40 | n/a |  |
|  | 12 | RA,5 | RA,5 | RA, IR,10 | RA, RF, IR,15 | RA, RF, IR,15 | RA, RF, HP, ND, IR, OM CY, 40 | n/a |

## Acknowledgments

We thank the animal caretakers, Rachel LaCasse, Danielle Adney, Thomas A. Bowden, Anita Mora, Robert Fischer, Myndi Holbrook, Ilona Rissanen, Emmie de Wit, and Trenton Bushmaker for their assistance during the study.

## Author contributions

N.v.D., T.L., S.C.G. and V.J.M. designed the study; N.v.D., V.A.A., R.F., J.E.S., E.H., A.O., J.L., P.W.H., K.C., G.S, and V.J.M. acquired, analyzed and interpreted the data; N.v.D. and V.J.M. wrote the manuscript. All authors have approved the submitted version.

## Data and materials availability

If data are in an archive, include the accession number or a placeholder for it. Here also include any materials that must be obtained through an MTA. Acknowledgments follow the references and notes but are not numbered.

